# Accelerating loss of resilience in Bornean rainforests

**DOI:** 10.1101/2025.07.21.665904

**Authors:** Shujie Chang, Neel P. Le Penru, Robert M. Ewers

**Affiliations:** Georgina Mace Centre for the Living Planet, Department of Life Sciences, Imperial College London, Ascot, UK; Dyson School of Design Engineering, Imperial College London, London, UK; Department of Zoology, University of Cambridge, Cambridge, UK; University Museum of Zoology, University of Cambridge, Cambridge, UK

**Author notes:** Corresponding authors: Shujie Chang, Neel P. Le Penru.

## Abstract

Climate change and human activity are eroding tropical rainforest resilience, risking tipping points of mass forest dieback towards savannah-like or treeless states. Here, we extend a close analysis of tipping-point early-warning signals in satellite-sensed vegetation data from the Amazon to Borneo, revealing resilience loss via critical slowing down of recovery from perturbation. Specifically, we examine spatiotemporal patterns in rising lag-1 autocorrelation and variance to assess whether, when and where resilience loss accelerated, and how these patterns relate to climate, human activity and animal biodiversity. We find that Bornean rainforests lost resilience over 1991–2016, with a marked acceleration from a breakpoint in 2004. Resilience loss accelerated earlier in subregions with higher human footprint, and was more severe in drier, more human-impacted subregions before the breakpoint. Post-breakpoint, resilience loss was stronger in wetter and less biodiverse areas. We also uncover the apparent reallocation of post-breakpoint resilience loss to areas that had previously appeared more resistant. Climate and bird and mammal richness appear to be most strongly correlated with this pattern. Our results provide enhanced evidence of Bornean rainforest resilience loss and its potential drivers, underscoring the urgent need for ecological conservation and climate change mitigation in Borneo and worldwide.

## Introduction

Tropical rainforests support over two-thirds of the world’s terrestrial biodiversity and provide critical functions in climate regulation, water cycling, and carbon sequestration^1,2^. Over recent decades, tropical regions including Borneo have experienced severe forest loss due to drought, fire, deforestation, and other pressures^3–5^. Worryingly, climate and land-use change may be driving tropical forests towards tipping points involving mass forest dieback, resulting in alternative savannah-like or treeless stable states^6,7^. This is alarming as tipping points can be especially difficult to reverse and predict from gradual trends in the mean state of the system alone^8,9^. Here, we build on work using satellite data to track resilience-based early-warning signals of tipping points in global vegetation^10–12^, to more closely examine trends in Bornean rainforest resilience loss and its possible drivers.

Tipping points are qualitative changes to the state of a system in response to small perturbations or increments in conditions^13^. Though various mechanisms may cause them^14^, tipping points are commonly associated with fold (a.k.a. saddle-node or catastrophic) bifurcations in dynamical systems, wherein stabilising negative feedback breaks down, triggering an abrupt shift to an alternative stable state driven by positive feedback^9,15^. As a system approaches a fold bifurcation, its dominant eigenvalue approaches zero from below and the system recovers more slowly from perturbations – so-called ‘critical slowing down’^16–18^. This slowing down reflects resilience loss, under the broad definition of resilience as the capacity to withstand disturbances, which bridges Holling’s canonical notions of ecological resilience (how much disturbance a system can withstand without changing its function) and engineering resilience (recovery rate from perturbation)^19,20^.

Crucially, critical slowing down causes systems to exhibit longer and larger temporal fluctuations about their current mean state, typically resulting in rising lag-1 autocorrelation (AC(1)) and variance, two classical early-warning signals of resilience loss toward a tipping point^21–23^. Other early-warning signals exist (for overviews see ref.^14,24^), but those based on generic statistical properties remain prominent and have been observed in long timeseries or along spatial stress/disturbance gradients in many natural systems approaching transitions. Examples range from yeast population collapses^25^ and lake eutrophication^26^ to the loss of major Earth-system elements such as the Greenland ice sheet^27^ and the Atlantic Meridional Overturning Circulation^8^.

Satellite data has proven invaluable for tracking early-warning signals in vegetation over decades at high resolution^11,12^, revealing worrying trends in global forest resilience loss. However, more detailed analyses of these trends and their drivers have focused primarily on the Amazon. In particular, Boulton et al.^10^ revealed a marked loss of Amazon rainforest resilience from the early 2000s, with faster loss in regions with lower mean annual precipitation and greater proximity to human activity. Furthermore, possible internal ecological drivers, particularly biodiversity, have received less attention. Although studies have demonstrated biodiversity’s importance to ecosystem resilience, most empirical analyses have focused on community-level processes and interactions^28–30^. Little work has empirically explored how biodiversity relates to nonlinear dynamical system models of complex ecosystems, despite longstanding deliberation on how biodiversity relates to ecosystem stability (e.g., ref.^31–33^).

Here, we extend the methods used by Boulton et al.^10^ to study rainforest resilience trends across Borneo. We measure the AC(1) and variance of satellite-sensed vegetation data and use breakpoint regression to examine whether and when trends in resilience-based early-warning signals changed. We also explore how climate, human activity and animal biodiversity relate to resilience change before versus after the breakpoint. Our analysis provides more detailed insight into Bornean rainforest resilience loss and its potential causes. Furthermore, we expand the toolkit for interpreting early-warning signals of critical transitions to quantify the timing and magnitude of accelerating resilience loss.

## Results

### Resilience change

We quantified the change in resilience of rainforest across Borneo from 1991 to 2016, focusing on 475 grid cells (0.25° × 0.25°) identified as intact rainforest, based on high forest cover (> 80% evergreen broadleaf cover) and the absence of human land use, as in previous work^10^ (Methods). We calculated two early-warning signals of a critical transition, AC(1) and variance, in sliding windows on detrended vegetation optical depth (VOD) time series (Methods). As AC(1) is generally a more robust indicator than variance^34,35^, we focused primarily on it in our analyses, and used variance as a supplementary metric to confirm that patterns in AC(1) likely stem from changes in ecosystem resilience^36^ (Methods). We then fitted a Bayesian breakpoint regression to the AC(1) time series and calculated the following three metrics of resilience change: (i) the consistency of resilience loss, measured by Kendall’s *τ* for the full, pre-breakpoint and post-breakpoint segments; (ii) the timing of the breakpoint; (iii) the magnitude of resilience loss, quantified using the posterior mean slopes of the pre- and post-breakpoint periods.

Throughout the study period, there was an upward trend in the mean AC(1) time series (Fig. 1; Kendall’s *τ* = 0.719, *p* < 0.001), indicating a consistent reduction in rainforest resilience. A breakpoint occurred around March 2004 (95% Credible Interval (CrI): July 2003–November 2004), after which the long-term upward trend became more pronounced (pre-breakpoint: Kendall’s *τ* = 0.387, *p* = 0.068; slope = 0.008 yr^−1^ [95% CrI: 0.006–0.010]; post-breakpoint: Kendall’s *τ* = 0.740, *p* < 0.001; slope = 0.032 yr^−1^ [95% CrI: 0.029–0.034]; Fig. 1). Model comparison showed that including a breakpoint provided a better description of the trend than a model without a breakpoint (leave-one-out information criterion, ΔLOOIC = −156.1). The timing of the breakpoint fell within a 5-year VOD window that was potentially influenced by the additions of the AMSR-E and WindSat sensors, but it lies towards the end of this period and does not fully overlap with it (Methods; Fig. S1). The mean VOD AC(1) also increased during a period without sensor changes, and sometimes decreased during periods with them, suggesting sensor changes are unlikely the main influence on AC(1) trends. Analysis of VOD variance also revealed an upward trend for the full and post-breakpoint time series (Fig. S2). AC(1) and variance were positively correlated (full period: Spearman’s *ρ* = 0.459; post-breakpoint: Spearman’s *ρ* = 0.533; both *p* < 0.001), indicating that the changes in these indicators most likely reflect actual resilience loss (i.e., slowing down of fluctuations^36^).

**Fig.1.**
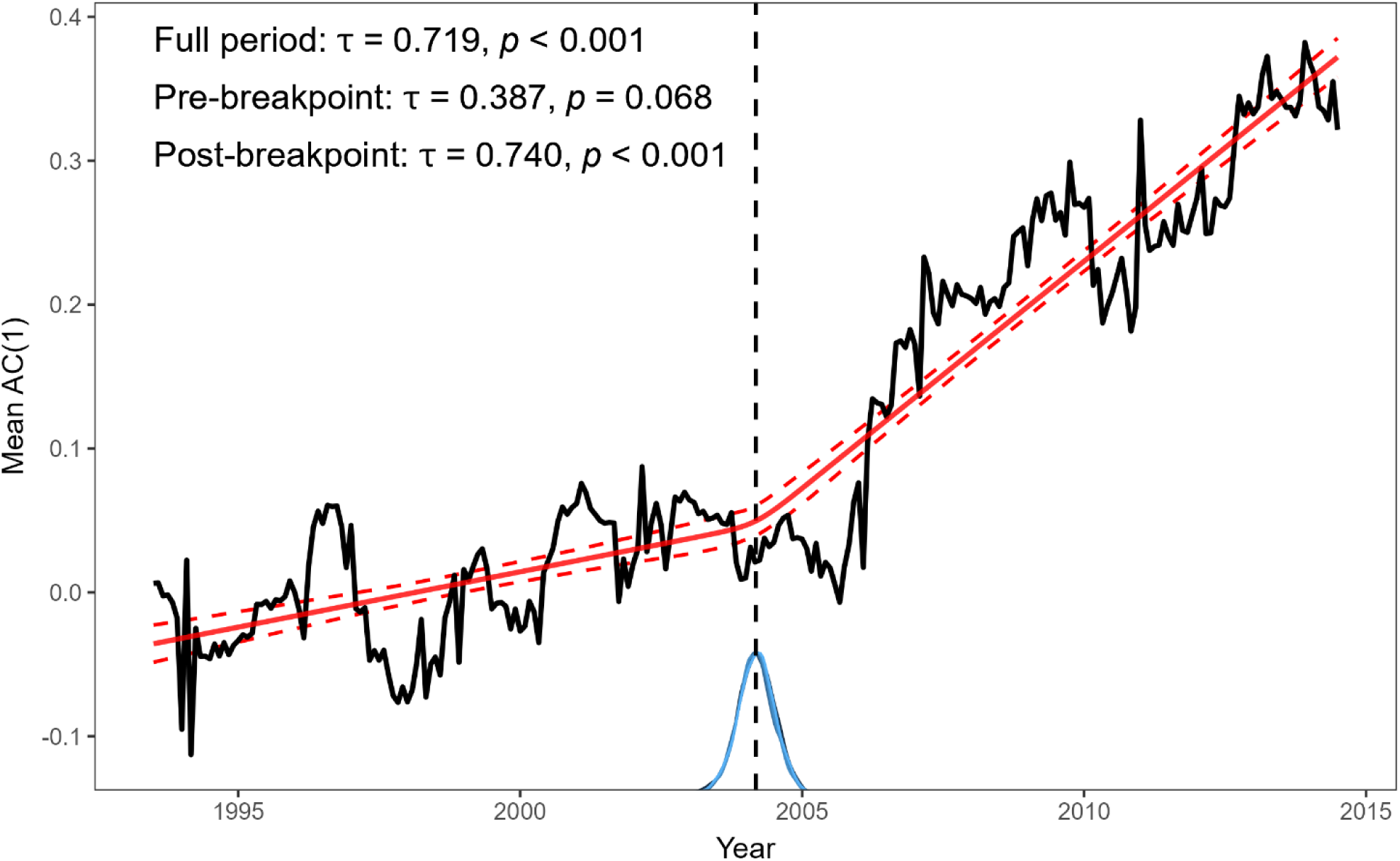
Overall resilience changes in Bornean rainforest. Mean VOD AC(1) time series of the study area from July 1993 to July 2014, calculated on 5-year monthly sliding windows and plotted at the midpoint of each window. The red line represents the posterior mean from a Bayesian breakpoint regression; the red dashed lines indicate the 95% credible interval based on 50,000 posterior draws (10,000 iterations × 5 chains). The vertical dashed line indicates the posterior mean breakpoint, with the blue curve below showing its posterior distribution. Kendall’s *τ* and associated *p*-values for the full record, pre-breakpoint, and post-breakpoint periods are reported at top left.

Across Borneo, individual grid cells exhibited breakpoints around the same time as that observed for the entire study area (mean across grid cells: February 2004; SD: 3.78 yr; Fig. 2a). Before the breakpoint, AC(1) increased in 52% of grid cells and across all grid cells the average Kendall’s *τ* was 0.027 (SD: 0.47; Fig. 2b). After the breakpoint, AC(1) increased in 65% of grid cells and the average upward trend was more pronounced, with a mean Kendall’s *τ* of 0.19 (SD: 0.47; Fig. 2c). Similarly, 51% of grid cells exhibited a positive slope pre-breakpoint, rising to 67% post-breakpoint (Fig. 2b–e). Slope values were also strongly correlated with Kendall’s *τ* (Fig. 2b–e; Table S1).

**Fig. 2.**
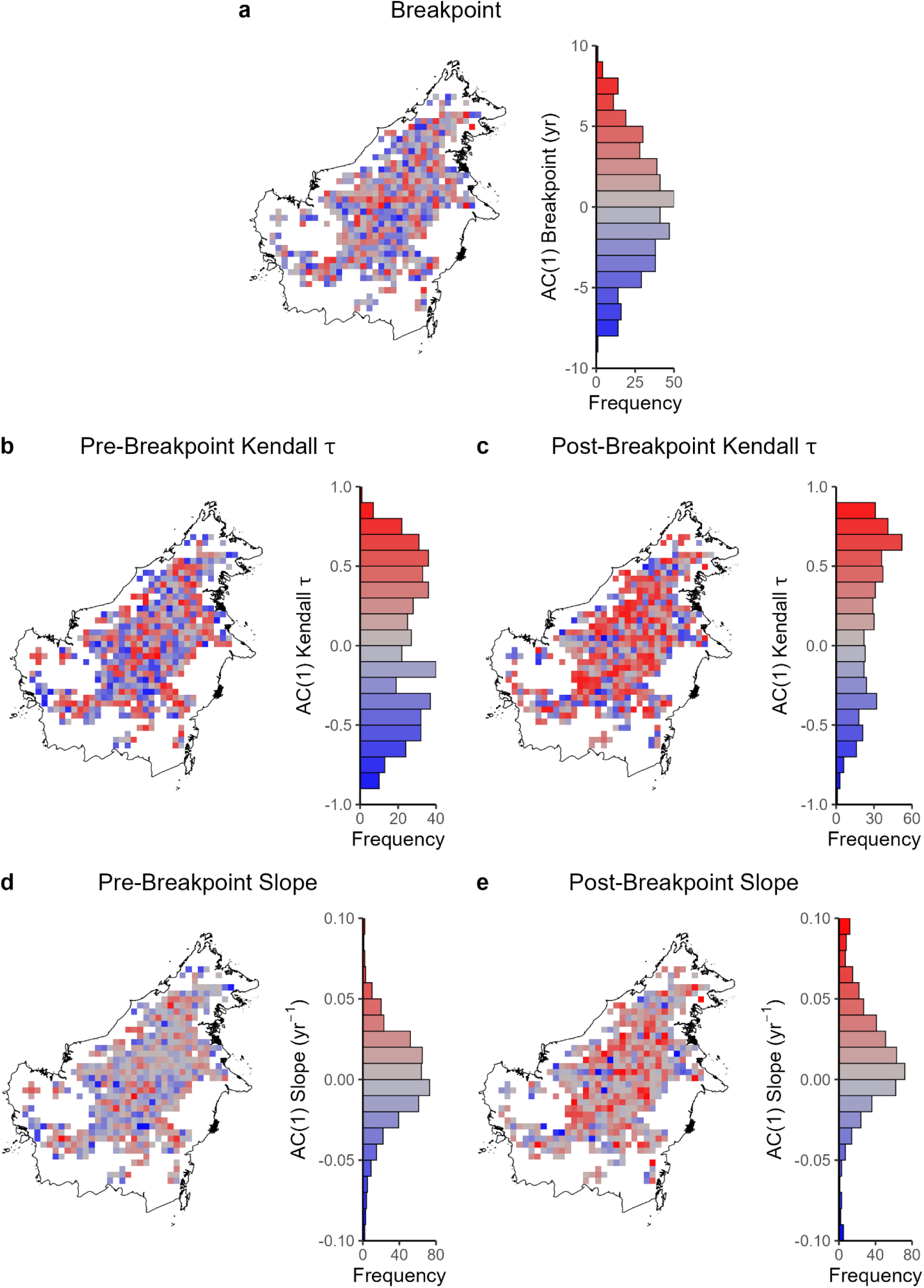
Spatial patterns of vegetation resilience changes in Bornean rainforest. **(a)** Map derived from the AC(1) time series for each grid cell. The vertical histogram serves as both a colour legend and the frequency distribution of map values. Colours encode the deviation of each cell’s breakpoint from the study-wide breakpoint (March 2004; positive values show later breakpoints). **(b–e)** Same layout as **(a)**, but for **(b)** pre-breakpoint Kendall’s *τ*, **(c)** post-breakpoint Kendall’s *τ*, **(d)** pre-breakpoint posterior mean slope (yr^−1^), **(e)** post-breakpoint posterior mean slope (yr^−1^). Note, Bayesian quantities in **(a)**, **(d)** and **(e)** are posterior means computed from 25,000 posterior draws (5,000 iterations × 5 chains). For visualisation, in **(d)** and **(e)**, values outside the plotting range are squished to the limits.

Breakpoint timing was only weakly correlated to resilience loss before and after the breakpoint (Table S1). Notably, the consistency and magnitude of resilience loss in individual grid cells were inversely associated between pre- and post-breakpoint periods (Fig. 2b–e; Table S1), suggesting that areas that lost more resilience before the breakpoint went on to lose less after it, and vice versa.

These results were broadly robust to varying broadleaf forest cover thresholds used to determine the study area (Figs. S3, S4). For the overall trend, the AC(1) time series showed a similar upward trend across thresholds (60%: Kendall’s *τ* = 0.71; 90%: Kendall’s *τ* = 0.709; both *p* < 0.001; Figs. S3a, S4a). The lower 60% threshold produced a broader breakpoint posterior with an earlier main peak, suggesting overall resilience loss in forest including less intact areas may have accelerated from December 2000 (95% CrI: March 1999–April 2004; Fig. S3a). Increasing to 90% broadleaf cover yielded little change relative to the 80% baseline (July 2004, 95% CrI: December 2003–February 2005; Fig. S4a). Spatially, the distribution of breakpoint timing and of pre- and post-breakpoint resilience changes (Kendall’s *τ* and slope) remained similar (Figs. S3b–f, S4b–f). The results were also robust to different AC(1) window lengths (Fig. S5).

For comparison to the Amazon, we recalculated the VOD AC(1) result in Boulton et al.^10^ using our processing pipeline (Methods). The AC(1) time series for the Amazon still shows an increasing post-breakpoint trend (Kendall’s *τ* = 0.556, *p* < 0.001; slope = 0.014 yr^−1^ [95% CrI: 0.012–0.016]), with breakpoint timing remaining around early 2000 (Boulton et al.^37^: January 2000 [95% CrI: November 1999–March 2000]; this study: March 2000 [95% CrI: December 1999–July 2000]; Fig. S6a). Whereas Borneo lost resilience throughout the whole study period, the Amazon showed overall gains in resilience (declining AC(1)) before its breakpoint. Furthermore, 82% of Amazon grid cells exhibited breakpoints later than that for the overall area (Fig. S6b) and breakpoint timing at the local scale was more strongly and positively correlated with pre-breakpoint resilience loss than in Borneo (Table S1). Similarly to Borneo, the consistency and magnitude of resilience change were inversely related between the between pre- and post-breakpoint periods (Table S1).

### Factors correlated with resilience change

To explore how climate, human activity and biodiversity relate to the resilience of Bornean rainforests, we quantified correlations between our resilience metrics and four factors: mean annual precipitation, mean annual temperature, human footprint and vertebrate species richness (amphibians, birds, mammals, and reptiles; Methods). Pairwise correlations among factors were weak (Fig. S7). Building on previous work^10,12^, we examined sliding bands of grid cells to aggregate those with similar values of each factor. This procedure reduces noise from other factors and reveals resilience change across subregions along the gradient of each factor (Methods).

Across all factors, breakpoint timing showed little relationship to these gradients, except for human footprint (Fig. 3a–d). The consistency and magnitude of resilience loss were generally higher after the breakpoint (Fig. 3e–l). Areas with lower precipitation had slightly earlier breakpoints (Fig. 3a) and were associated with more consistent and stronger resilience loss before the breakpoint (blue points; Fig. 3e,i), but wetter regions generally experienced more resilience loss after the breakpoint (red points; Fig. 3e,i). Breakpoints tended to occur earlier in hotter areas (Fig. 3b), which also experienced less resilience loss pre-breakpoint (blue points; Fig. 3f,j). However, post-breakpoint associations between temperature and the consistency and magnitude of resilience change were mixed (red points; Fig. 3f,j). Interestingly, along both the precipitation and temperature gradients, pre- and post-breakpoint values of consistency and magnitude were negatively correlated (Fig. 3e,f,i,j; Table. S2). Hence, the relationship between climate and resilience seemed to invert post-breakpoint. As human footprint increased up to a value of 2 units, areas exhibited earlier breakpoints, but the relationship between human footprint and breakpoint timing was relatively flat beyond this (Fig. 3c). Areas with more intensive human activity showed less consistent pre-breakpoint resilience loss (blue points; Fig. 3g) and lower magnitude resilience loss pre- and post-breakpoint (Fig. 3k).

**Fig. 3.**
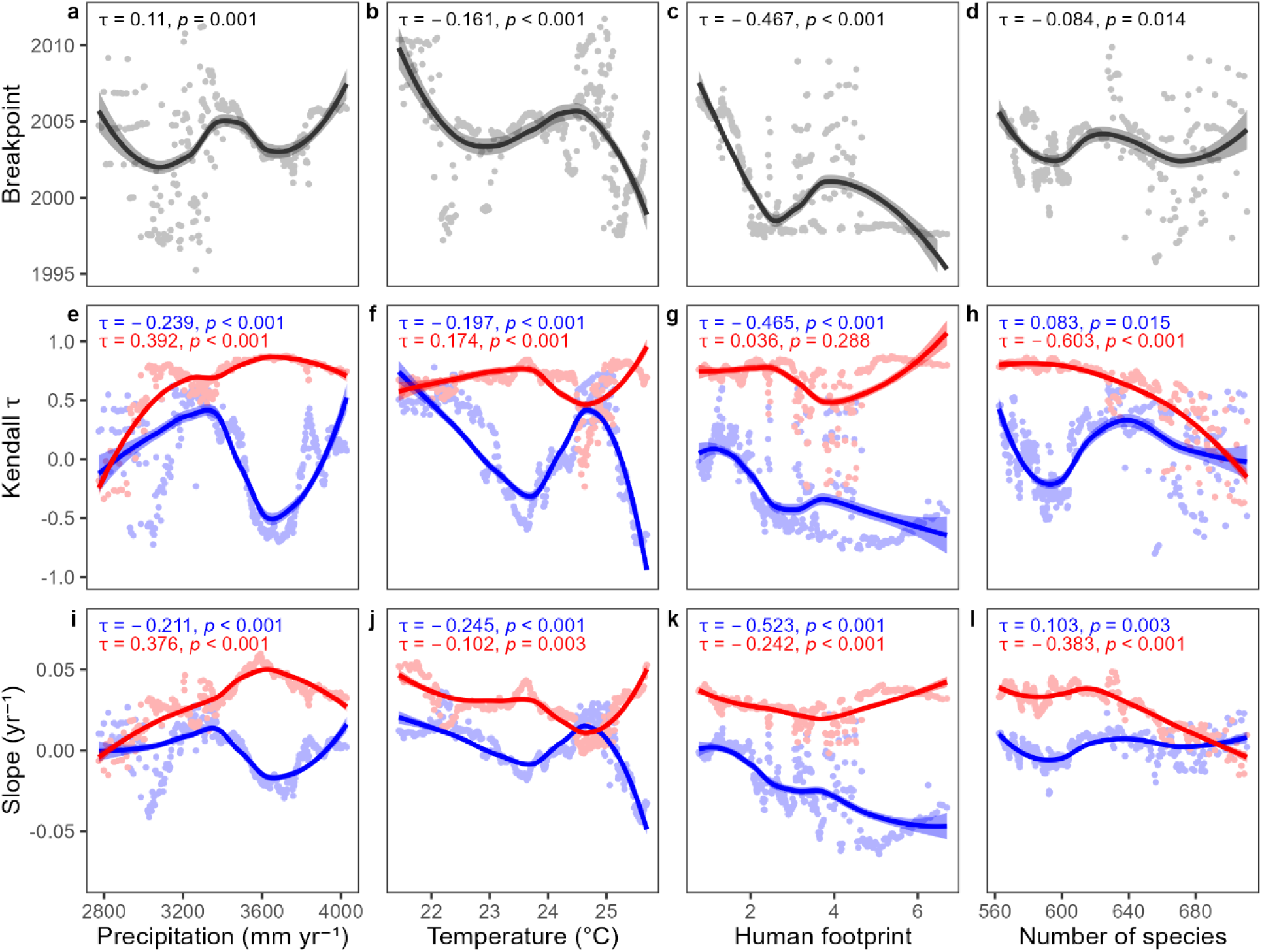
Relationship between vegetation resilience and potential influencing factors (sliding band size = 100 cells). Rows correspond to different resilience metrics and columns represent different factors. **(a, e, i)** Correlation between precipitation and the following resilience metrics calculated on sliding bands of grid cells along a precipitation gradient (Methods): **(a)** posterior mean breakpoint from Bayesian breakpoint regressions; **(e)** Kendall’s *τ* (blue: pre-breakpoint; red: post-breakpoint); and **(i)** posterior mean slope (blue: pre-breakpoint; red: post-breakpoint). The correlations between the resilience metrics and potential influencing factors are estimated using Kendall’s *τ*. the Kendall’s *τ* and associated *p*-values are reported at top of each panel. **(b–d, f–h, j–l)** The same as **(a, e, i)**, but for **(b, f, j)** temperature, **(c, g, k)** human footprint and **(d, h, l)** animal species richness, respectively. In each panel, the values are plotted at the midpoint of each band.

For animal biodiversity, breakpoint timing (Fig. 3d) and pre-breakpoint resilience loss (blue points; Fig. 3h,l) had inconsistent relationships to overall species richness. However, post-breakpoint resilience loss was notably less consistent and pronounced in areas of high biodiversity (red points, Fig. 3h,l). Individual taxa were intercorrelated to varying degrees (Fig. S8), but the overall patterns (Fig. 3d,h,l) were broadly consistent across groups (Fig. S9). The resilience change metrics along the overall species richness gradient had weak or non-significant correlations between their pre- and post-breakpoint values (Table S2). However, birds and mammals exhibited strong inverse correlations in their pre- and post-breakpoint values of Kendall’s *τ* and slope, whereas amphibians showed positive pre- versus post-breakpoint correlations (Table S2). The above patterns across all factors and taxonomic groups were robust to the choice of band size in sliding-band analyses (Figs. S10, S11).

## Discussion

We found that, similarly to the Amazon^10^, Bornean rainforests have been losing resilience. However, unlike the Amazon, Bornean rainforests lost resilience throughout the study period, although the rate at which resilience was lost has notably accelerated from the early 2000s. Overall, the long-term average rate at which resilience is being lost is higher in Borneo than it is in the Amazon. This could mean tipping points are more imminent in Southeast Asia than in the Amazon, though we emphasise that the timing of tipping points depends on many factors and remains notoriously hard to predict. In addition, differences in breakpoint timing between these two regions (Figs. 1, 2, S6) suggest that pan-tropical resilience loss is likely driven by regional factors and/or local responses to global climatic trends, rather than a single global event.

Differences in local (individual grid-cell) resilience dynamics may offer further insight into the mechanisms underlying resilience loss. In Borneo, local breakpoints were nearly synchronous with the landscape-scale (spatially-aggregated) breakpoint, and were only weakly correlated with the degree of resilience loss before the breakpoint. This pattern may indicate that resilience loss is dominated by broad-scale external forcing that pushes many areas towards a transition at roughly the same time. In the Amazon, local breakpoints mainly lagged behind the landscape-scale breakpoint and were more strongly correlated with pre-breakpoint resilience loss. This pattern may arise from gradual resilience loss in subregions of the Amazon that exert an outsized influence over other large areas. For instance, deforestation in upwind regions reduces evapotranspiration, with severe consequences for downwind forests that lose an important share of their precipitation as a result^38^. Together, these patterns suggest that tropical forests may respond in different ways to drivers operating at different scales and to different stages of resilience loss.

Although Borneo lost resilience overall and within most subregions, a number of grid cells in Borneo did not, and even appear to have gained it. Further, in both Borneo and the Amazon, places that lost more resilience before the breakpoint tended to lose less—or even gain resilience— afterward, and vice versa. While further work is needed to understand this pattern, it reveals that landscape-wide resilience loss arises from a mosaic of local patterns and factors that can be asynchronous and interact in various elaborate ways, including this apparent reallocation of resilience between subregions. More generally, this finding underscores the complexity and scale-dependent nature of ecosystem resilience^39–41^.

Across Borneo, the onset of accelerated resilience loss tended to occur earlier in drier and hotter regions. Such areas have been shown to exhibit lower functional diversity, declining functional redundancy and heightened vulnerability^42,43^. Grid cells with lower precipitation lost more resilience before the breakpoint, broadly comparable to prior findings^6,10,12^. However, after the breakpoint, wetter regions experienced more pronounced resilience loss, perhaps because high precipitation can become a stressor once forest condition is compromised (e.g., via waterlogging and root hypoxia, heightened pathogen and/or pest pressures^44–46^). By contrast, no clear patterns of pre- and post-breakpoint resilience loss emerged along the temperature gradient, perhaps because temperatures largely fell within a suitable range and were not a key stressor in Borneo.

The reversal of patterns between climate and resilience after the breakpoint may account for the apparent spatial reallocation of resilience post-breakpoint described earlier, for example, via landscape-scale water balance reorganization driven by moisture-recycling^38,47^. Such asynchrony in local stress may produce insurance-like compensatory dynamics that redistribute resilience across the landscape, helping to maintain ecosystem functioning. However, further work is needed to understand whether and why this might be the case, and to explore whether these findings are an artefact of the breakpoint method and treating drivers as time-invariant.

Regarding human activity, areas with higher human footprint tended to reach the breakpoint earlier (although this response rapidly plateaus), consistent with anthropogenic disruption of key ecological processes and networks, such as seed dispersal and pollination, which can increase vulnerability to disturbances^48,49^. Human-induced fires and deforestation can also produce a drier microclimate that makes it easier for subsequent fires to start and spread^6,38^. In contrast to findings from the Amazon^10^, we observed that areas with greater human footprint were associated with less pronounced loss of resilience overall. This could be indicative of positive human influences, such as reforestation and forest restoration efforts, but this interpretation requires further validation.

For biodiversity, we found that areas with higher species richness lost less resilience after the breakpoint. This pattern held for overall animal biodiversity and, broadly, within individual animal taxa. Like climate, biodiversity also appeared to modulate the spatial reallocation of resilience loss after the breakpoint, but this effect was clearest for birds and mammals, which showed strong reversals in the pre- and post-breakpoint relationships between richness and resilience. This pattern is consistent with an asynchrony-based insurance mechanism, whereby higher richness increases response diversity and the potential for asynchronous species dynamics across space and time, buffering shocks and shaping spatial heterogeneity in resilience change^50,50,51^. By contrast, amphibians displayed more similar pre- and post-breakpoint relationships to resilience, while reptiles showed less consistent patterns.

One explanation is that biodiversity simply tracks resilience rather than reinforcing it. Short-term resilience loss may not immediately trigger a regime shift, but it can still manifest in subtler ecosystem impacts (e.g. leaf wilting, fruit scarcity) that prompt animals to move towards more resilient areas in search of resources. Under this scenario, spatial variation in richness would emerge from differential movement rather than from biodiversity itself stabilising ecosystem dynamics. The IUCN biodiversity layers used here are aggregated over decades and lack population-level and movement resolution, which limits our ability to distinguish between these pathways. The presence or absence of a reallocation pattern may also result from variation in regional species-pool sizes, as the relatively small species pools of amphibians and reptiles may limit our ability to detect such richness effects in our spatially aggregated data. Disentangling these mechanisms will require community-level studies that couple resilience dynamics with interaction-network structure, functional roles, and high-resolution monitoring of populations and movement.

By introducing two additional metrics (breakpoint and slope) of resilience change, we captured more detail on the correlates and potential drivers of resilience loss in Bornean rainforest. We emphasize the analysis is correlational, however; further investigation is needed to determine whether these relationships are causal and exactly how they operate. Moreover, ecosystem dynamics and resilience are complex, involving potentially many factors beyond those considered here, as well as indirect or multivariate effects^14,52,53^. However, those we examined are in line with previous work and are considered the predominant global and local factors impacting rainforests worldwide^6,7,10,12,50,51,54^.

In addition, early-warning signals derived from critical slowing down cannot accurately predict the type of impending critical transition or when it might occur, may produce false alarms (e.g., when the sources of noise or random disturbances change^35,55^), and critical transitions may occur without critical slowing down^56–58^. However, we substantially reduced the likelihood of false or missed alarms by examining both AC(1) and variance. Furthermore, these early-warning signals remain useful in suggesting increased forest instability and vulnerability to hazards, which could offer actionable guidance to Bornean conservation initiatives, such as the Heart of Borneo^59^. Integrating our results alongside more fine-grained biodiversity and ecosystem health monitoring, as well as with larger scale mechanistic models of rainforests, may help further overcome the above risks and produce a more detailed picture of Bornean rainforest health and resilience.

We found that long-term resilience loss in rainforests is a pan-tropical issue, but it appears to be shaped by spatial variations in biotic and abiotic factors. Our findings also indicate that regional variation in key global drivers of rainforest resilience loss (e.g., global warming) and local factors interplay to produce differences in the patterns and underlying mechanics of Bornean versus Amazonian rainforest resilience loss. Moreover, climate-driven asynchrony in local stress and biodiversity-driven asynchrony in responses may generate an insurance-like, compensatory reallocation of resilience across space and time. These insights may inform scientific guidance for the conservation and management of rainforests to avert tipping points of mass vegetation mortality. Safeguarding resilience will require cross-scale strategies that mitigate global climate change, preserve landscape connectivity and moisture-recycling pathways, maintain response diversity, and embed site-level actions within coordinated, adaptive portfolios.

## Methods

### Datasets and preprocessing

We defined the geographical extent of Borneo (7°N to 4°S, 108°E to 119°E) by integrating shapefiles of Malaysia, Brunei, and Indonesia from the GADM database version 4.1 (https://gadm.org/data.html, accessed 17 November 2023). To identify the intact rainforest of Borneo as our study area, we applied filters to the International Geosphere-Biosphere Programme (IGBP) Moderate Resolution Imaging Spectroradiometer (MODIS) land cover dataset MCD12C1 v061^60^. We selected grid cells where evergreen broadleaf forest cover was ≥ 80% in 2001 and excluded any cells containing human land use (built-up, croplands, or vegetation mosaics > 0%^10^).

To evaluate the resilience of intact Bornean rainforest, we used VOD data from the Vegetation Optical Depth Climate Archive^61^ (VODCA). Compared to optical remote sensing vegetation products, such as Normalized Difference Vegetation Index (NDVI), passive microwave-derived VOD has the advantages of high sensitivity to high biomass levels, and the capability to perform under cloud cover, making it particularly suitable for tropical forest analysis^62–64^. VOD products with shorter wavelengths are more sensitive to leaf moisture content and better reflect changes in vegetation resilience, but are less influenced by stem biomass and soil moisture^61,65,66^. The Ku-band of VODCA was selected because it is the shortest wavelength band of the dataset, while providing the longest time span of short wavelength products by combining multiple sensors^61,65^.

To explore factors that may affect resilience change, we considered three perspectives: climate, human activity, and biodiversity. For climate, we focused on precipitation and temperature, sourced from the monthly average precipitation data of the Climate Hazards Center InfraRed Precipitation with Station data (CHIRPS) v2.0 dataset^67^ and the fifth generation of European ReAnalysis (ERA5) Land monthly average 2 m air temperature dataset^68^, respectively. Impacts of human activity were assessed using human footprint data from the database described by Mu et al.^69^. To avoid potential interference between plant diversity and VOD-derived resilience, we focused on animal biodiversity when exploring the relationship between biodiversity and resilience. Specifically, we utilized animal species richness data from the International Union for Conservation of Nature (IUCN) Red List version 2022-2^70^, including four taxa groups (amphibians, birds, mammals, and reptiles).

Preliminary exploration of the data highlighted significant fluctuations in the first three years of VODCA data, which were also reported by Boulton et al.^10^. Therefore, we also restricted our analysis to 1991**–**2016. The land cover dataset was only available from 2001 and was used up to 2016 to match the study period. We adopted a 0.25° × 0.25° spatial resolution, matching that of the VOD data (the coarsest dataset we examined) and unified the data under the WGS 1984 geographic reference system. We then aggregated data with different collection frequencies to the required analysis scale and cropped all data into the study area using the package *terra*^71^ (v.1.8.54) from R^72^ (v. 4.5.0).

### Generation of resilience indicators

We used the VOD time series to estimate the resilience indicators, AC(1) and variance. We first applied the Seasonal and Trend decomposition using Loess (STL) to remove the long-term trends and seasonal fluctuations from the VOD data^73^, to obtain a stationary time series of fluctuations to the system state, mostly caused by small perturbations^74,75^. We used the *stl*() function to decompose the time series in each grid cell (using the ‘*periodic*’ option for seasonal windows), retaining the residuals for subsequent analysis^10^.

AC(1) was calculated by fitting an order 1 autoregressive model^22^:

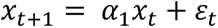

fit using ordinary least squares (OLS) with Gaussian random error *ɛ*_*t*_. This formula does not include an intercept because the model’s analysis was performed on detrended residuals with a mean of zero^22^.

We calculated resilience indicators over a 5-year (60-month) monthly sliding window, creating a new time series from July 1993 to July 2014. Importantly, to calculate Borneo-wide overall resilience indicators, we first spatially averaged the monthly VOD time series and then computed AC(1) and variance on the mean VOD series. This order of operations has proven more effective in reducing bias, especially that which arises from mixing sensors, compared to averaging resilience indicators computed for each grid cell^36^. To explore the spatial variation of resilience loss across Borneo, we calculated the resilience indicators for each grid cell following the same sliding window approach. This procedure yielded time series at two spatial scales—for entire Borneo and each pixel.

Previous work has shown that AC(1) is a more robust indicator when analysing resilience^34,35^, especially as variance is more susceptible to noise in ecosystem data and changes in environmental perturbations, hence we focus primarily on AC(1) in our analyses. However, Smith et al.^36^ recently highlighted that combining data from different sensors, such as VODCA, may also contribute to a rise in autocorrelation. They also point out that combined sensor data may bias AC(1) and variance toward negative correlation with one another, whereas actual resilience loss should yield increases in both AC(1) and variance^10,36^. Therefore, we mainly employ variance as a supplementary metric to validate that the increases in AC(1) are likely due to resilience loss. To further examine the potential influence of satellite sensor transitions on the VODCA products, we incorporated the timing of these shifts into the mean AC(1) series and compared these with the breakpoint (Fig. S1).

### Metrics of resilience indicators

We applied a Bayesian changepoint detection algorithm to reveal when trends in resilience changed, employing the package *mcp*^76^ (v.0.3.4). To assess whether a breakpoint was warranted, we compared a one-break segmented model to a no-break linear reference using Pareto-smoothed importance-sampling leave-one-out cross-validation (PSIS-LOO), with smaller ΔLOOIC values indicating better expected out-of-sample predictive fit^77^. Though the Kendall’s non-parametric rank correlation coefficient (*τ*) is used as a general indicator for evaluating trends in other parts of the study, in the analysis of trends in resilience it was specifically used to assess the consistency of changes in the AC(1) time series for the full, pre- and post-breakpoint periods. Kendall’s *τ* returns values of 1 for a consistently increasing series, −1 for a consistently decreasing series, and 0 when there is no overall trend. To specifically examine the magnitude of change in the rate of resilience loss around the breakpoint, we also extracted the posterior mean slopes of the linear segments before and after the inferred breakpoint from the Bayesian changepoint model. Within each segment, positive slopes indicate resilience loss, whereas negative slopes indicate resilience gain, with larger absolute values denoting a greater magnitude of change in the rate.

The *p*-value directly derived from the Kendall’s *τ* coefficient may be inaccurate due to its sensitivity to the autocorrelation characteristic inherent in time series data. Therefore, following prior work, we employed a phase randomization procedure to generate surrogate data to determine the significance of Kendall’s *τ*^10,22,78^. Specifically, we performed a Fourier transform on the AC(1) and variance time series, randomly permuted the phases, and then applied an inverse Fourier transform, thereby maintaining the autocorrelation and variance in the surrogate series^10,79^. We performed this 100,000 times to create a null model for assessing the significance of Kendall’s *τ*. The *p*-value was determined by calculating the proportion of surrogates that exhibit a higher Kendall’s *τ* value than that observed in the actual series.

Thess three metrics of resilience were applied to AC(1) time series at both spatial scales (Borneo-wide and per-pixel) to evaluate the resilience patterns. For variance, we only calculated Kendall’s *τ* of the overall time series, and used Spearman’s rank correlation coefficients (*ρ*) to quantify its association with AC(1), helping to distinguish genuine resilience loss from artefacts of multi-sensor assimilation (see ref.^36^). For each pixel, we calculated Spearman’s *ρ* for cross-metric and pre/post-breakpoint associations, including: (i) breakpoint timing vs. pre-break Kendall’s *τ*/slope—association between antecedent change and the onset of following change; (ii) breakpoint timing vs. post-breakpoint Kendall’s *τ*/slope—association between breakpoint and subsequent change; (iii) pre- vs. post-breakpoint values of Kendall’s *τ*/Slope—change across the breakpoint; (iv) Kendall’s *τ* vs. Slope—consistency of these two metrics.

### Factors correlated with resilience change

We used four factors—mean annual precipitation, mean annual temperature, human footprint and species richness—to explore potential relationships between vegetation resilience and climate, human activity, and biodiversity. We treated each factor as a constant, to focus on changes in resilience and standardize the analysis of all factors. Factors with time-series data, such as precipitation and temperature, were thus averaged over the study period for subsequent analysis, allowing us to focus solely on the spatial patterns in the factors and resilience change. We also calculated Spearman’s *ρ* to assess spatial correlations between the factors, given several factors had non-normal distributions and could have nonlinear relationships.

We explored the impact of various factors on resilience by ranking the values of each factor and applying a sliding band, where a fixed number of grid cells of 100 were used across the data range, shifting the band by one grid cell at each step. These bands were then used to spatially subset grid cells that fell within the range of factors’ values for each band. For each subset, we calculated the AC(1) from the VOD time series of the selected grid cells following the same approach used for the entirety of Borneo and estimated the three metrics of resilience change. Finally, we used Kendall’s *τ* to assess the correlation between the resilience metrics and factors.

### Robustness tests

To test the robustness of the results, we varied the evergreen broadleaf forest cover thresholds used to define the study area. Specifically, we estimated the AC(1) trend when the forest cover threshold is set to ≥ 60% and ≥ 90% (Figs. S3, S4). We also tested the robustness of the sliding window length used to calculate AC(1) by trying windows of 30, 90, and 120 months (Fig. S5). For exploration of the relationship between resilience loss pattern and factors, we also use sliding bands of 50 grid cells (Figs. S10, S11).

## Data Availability

The shapefiles for the countries are available from https://gadm.org/data.html. The VOD dataset is available from https://zenodo.org/record/2575599#.YHRE6RRKj7E. The MODIS Land Cover dataset is available from https://lpdaac.usgs.gov/products/mcd12c1v061. The CHIRPS precipitation dataset is available from https://data.chc.ucsb.edu/products/CHIRPS-2.0/global_monthly/tifs. The ERA5-Land temperature dataset is available from https://cds.climate.copernicus.eu/cdsapp#!/dataset/reanalysis-era5-land-monthly-means?tab=overview. The human footprint dataset is available from https://figshare.com/articles/figure/An_annual_global_terrestrial_Human_Footprint_dataset_from_2000_to_2018/16571064. The IUCN Red List Species Richness dataset is available from https://www.iucnredlist.org/resources/other-spatial-downloads.

## Acknowledgements

We thank Chris A. Boulton for helpful discussions as well as members of the Ewers group for feedback on earlier drafts, including David Orme, Vivienne P. Groner, Jacob Cook, Hollie Folkard-Tapp and Olivia Daniel. N.P.L.P. was supported by the Natural Environment Research Council (grant number NE/S007415/1) via the Science and Solutions for a Changing Planet DTP.

## Author contributions

The study was conceptualised by all authors. Data processing and statistical analyses were conducted by S.C. The project was supervised by R.M.E and N.P.L.P. The original draft was written by S.C. and N.P.L.P. All authors contributed to revising and editing the final manuscript.

## Supplementary Figures

**Fig. S1.**
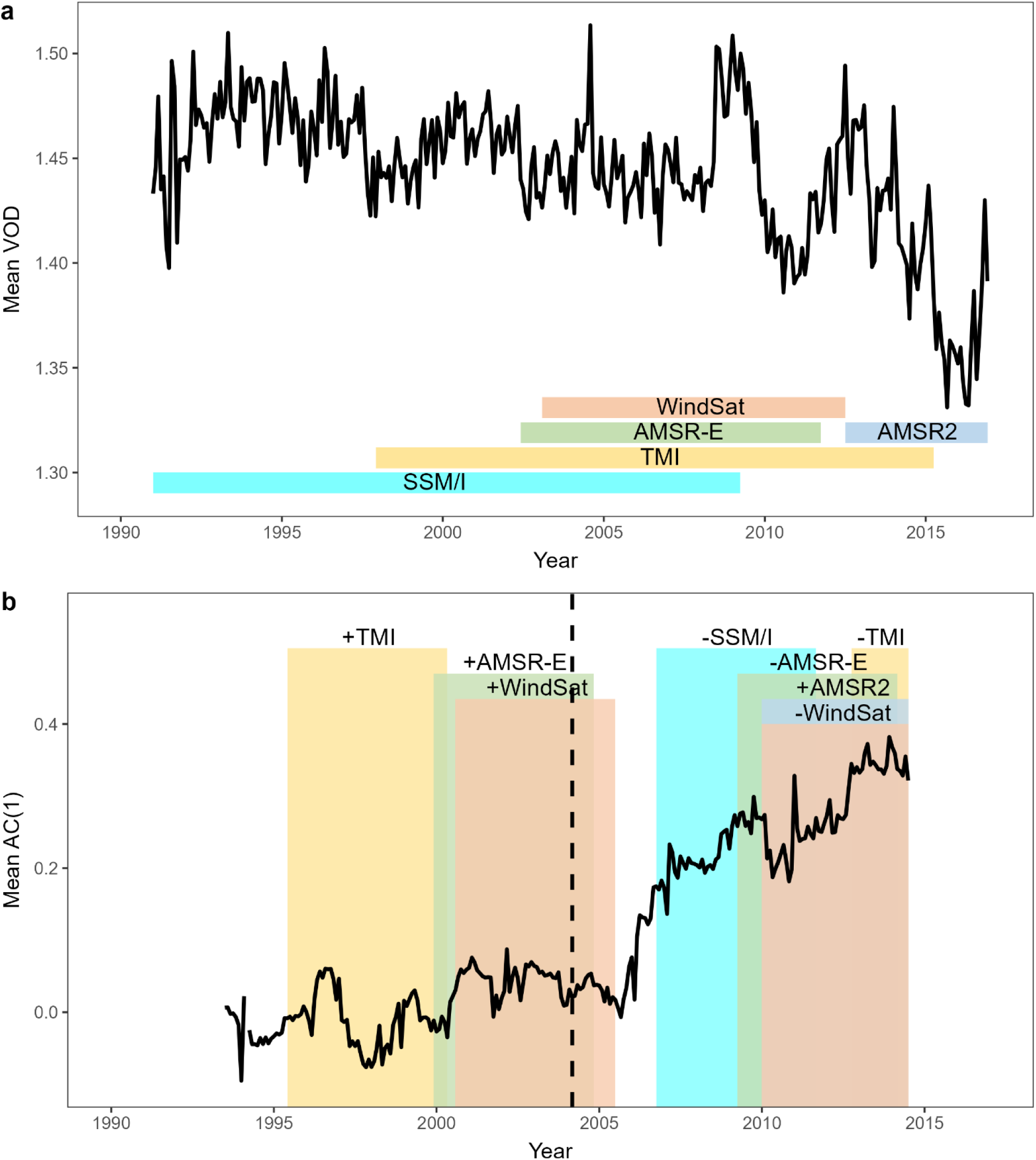
Relative time and impact of sensor shift on time series of VOD and AC(1). **(a)** Time series of monthly mean VOD spatially averaged over the study area, with the time span of each satellite sensor used to measure VOD shown in the bottom, adapted from Moesinger et al. (2020). **(b)** Mean AC(1) time series created from mean VOD time series above, calculated on 5-year sliding windows with monthly increments and plotted at the midpoint of each window. The coloured bands correspond to the addition (+) or removal (−) of sensors and highlight sections of the AC(1) time series that were calculated on windows of VOD data that feature those particular sensor changes. The vertical dotted line indicates the breakpoint of AC(1) time series.

**Fig. S2.**
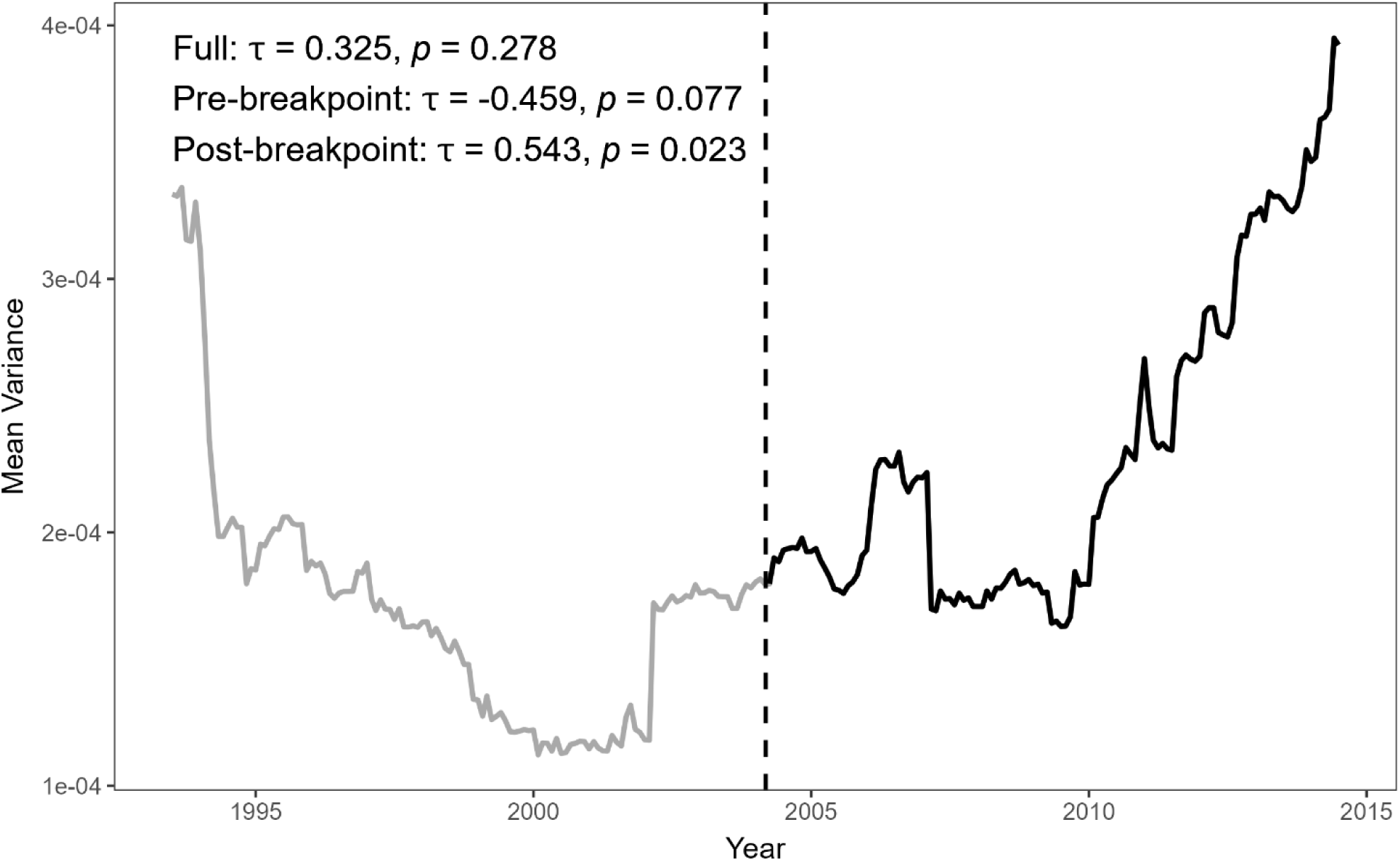
Changes in Borneo rainforest resilience since the 1990s and from the breakpoint as measured from variance of VOD. Mean VOD variance time series of the study area from July 1993 to July 2014 (compared to Fig 1). The vertical dotted line indicates the breakpoint determined from the AC(1) time series. Note that the variance values are plotted at the middle of each 5-year sliding window.

**Fig. S3.**
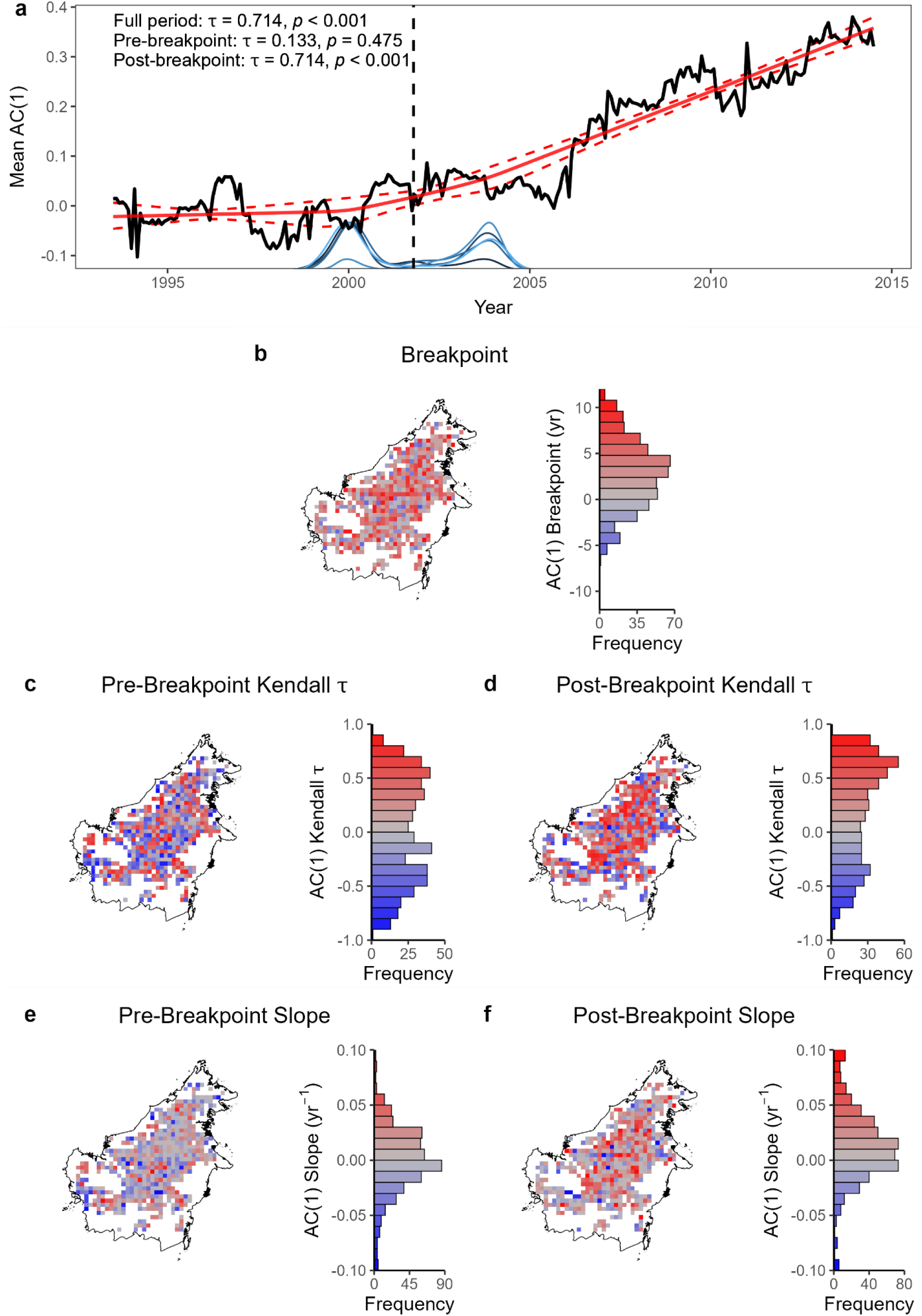
Overall trend and spatial patterns in resilience changes of Bornean rainforest when varying the threshold of evergreen broadleaf forest cover to 60% to determine study area. Recreation of Fig. 1 **(a)** and Fig. 2 **(b–f)** but using grid cells with forest cover ≥ 60%.

**Fig. S4.**
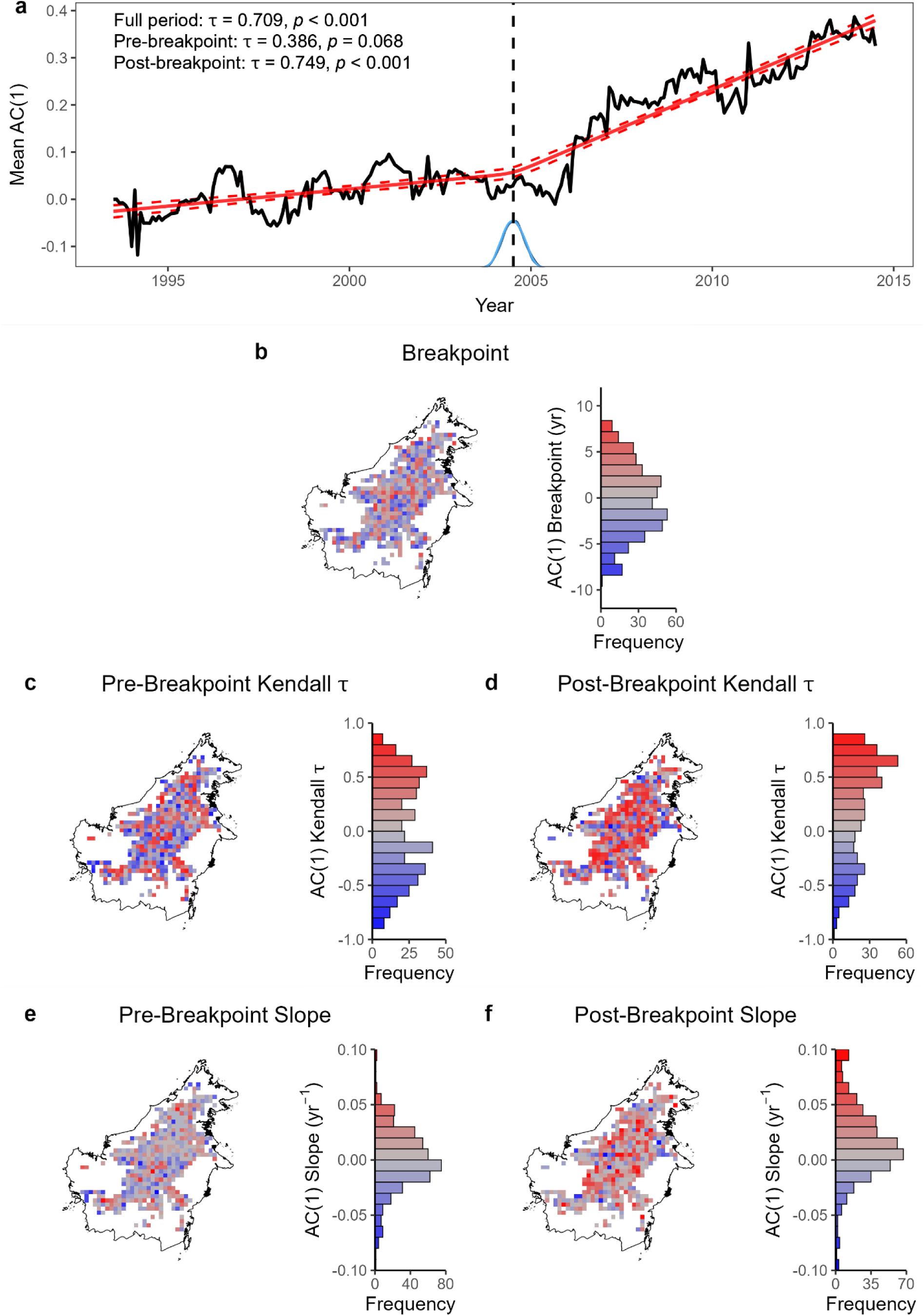
Overall trend and spatial patterns in resilience changes of Bornean rainforest when varying the threshold of evergreen broadleaf forest cover to 90% to determine study area. Recreation of Fig. 1 **(a)** and Fig. 2 **(b*–*f)** but using grid cells with forest cover ≥ 90%.

**Fig. S5.**
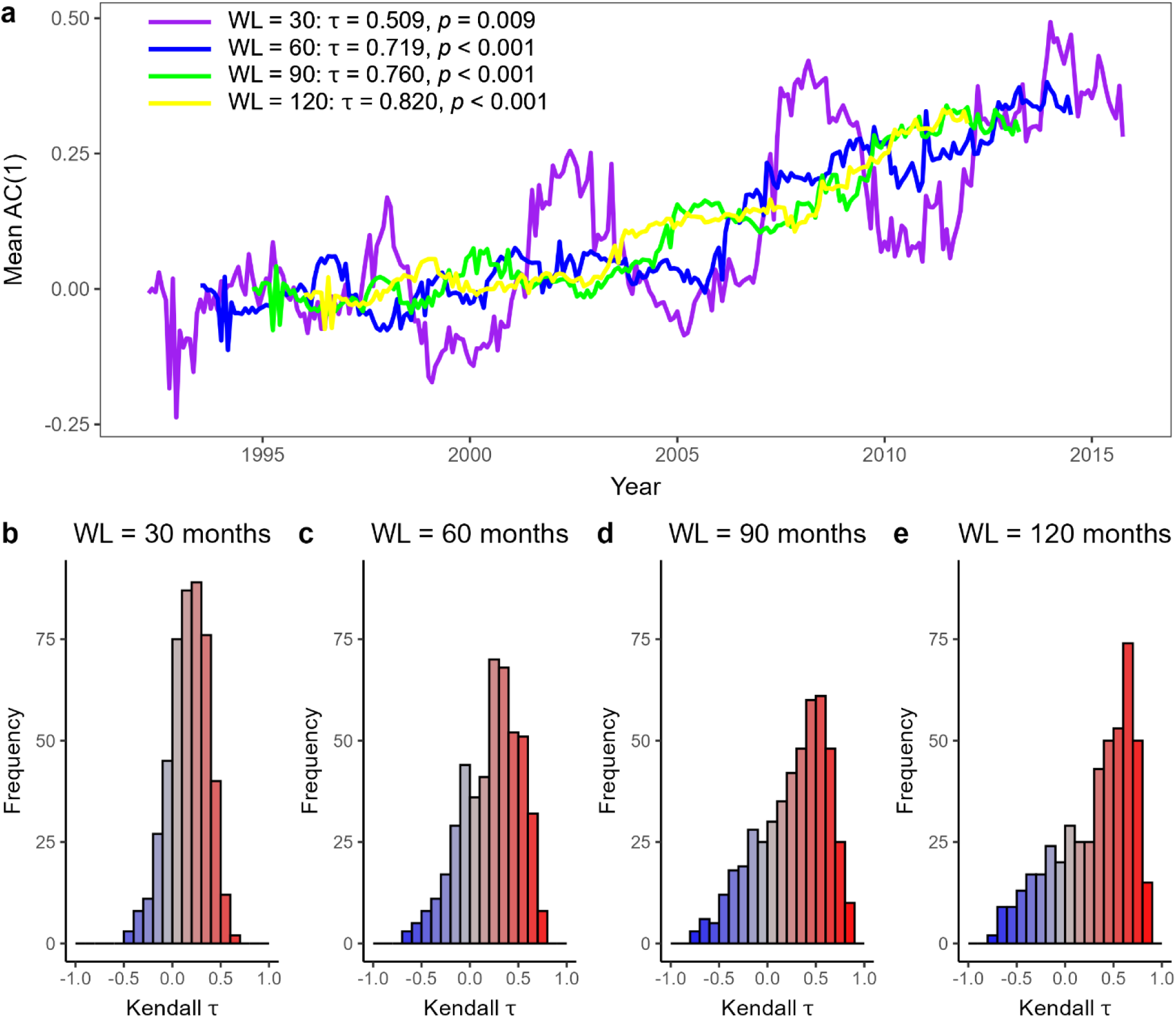
Robustness of window length to calculate VOD AC(1). **(a)** Mean AC(1) time series when varying the window lengths of calculating to 30 months, 60 months (as in Fig. 1), 90 months and 120 months. Note that the AC(1) values are plotted in the middle of the sliding window. **(b*–*e),** Histograms of grid cells’ Kendall’s *τ* values for these choices of AC(1) window lengths. WL = window length.

**Fig. S6.**
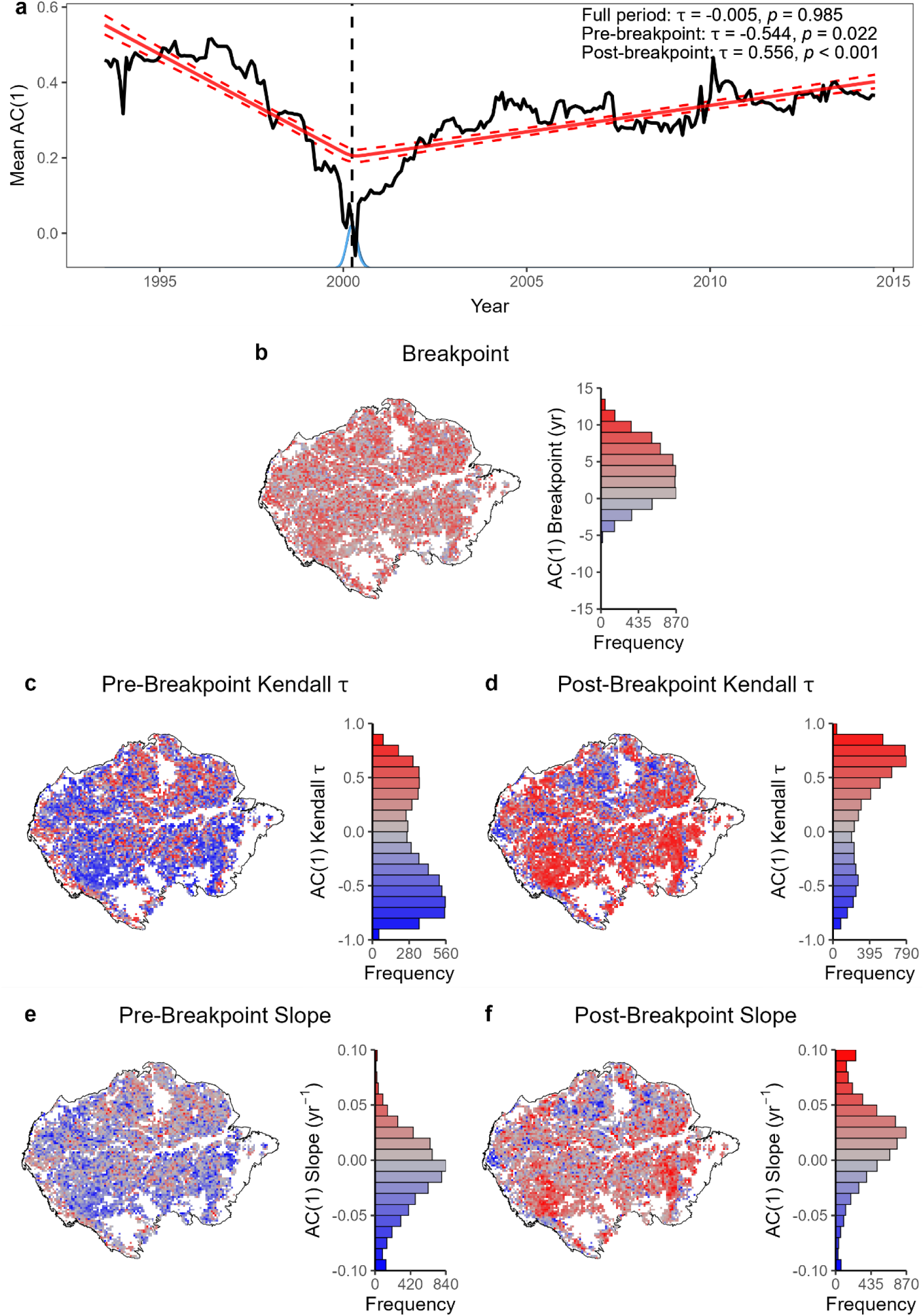
Overall trend and spatial patterns in resilience changes of Amazon rainforest, adapted from Boulton et al. (2022) with our processing pipeline. Compared to Fig. 1 **(a)** and Fig. 2 **(b*–*f)** but for the study area of the Amazon basin. Note that in our processing pipeline, VOD data are spatially averaged before calculating the overall mean AC(1) time series plotted in **(a)**.

**Fig. S7.**
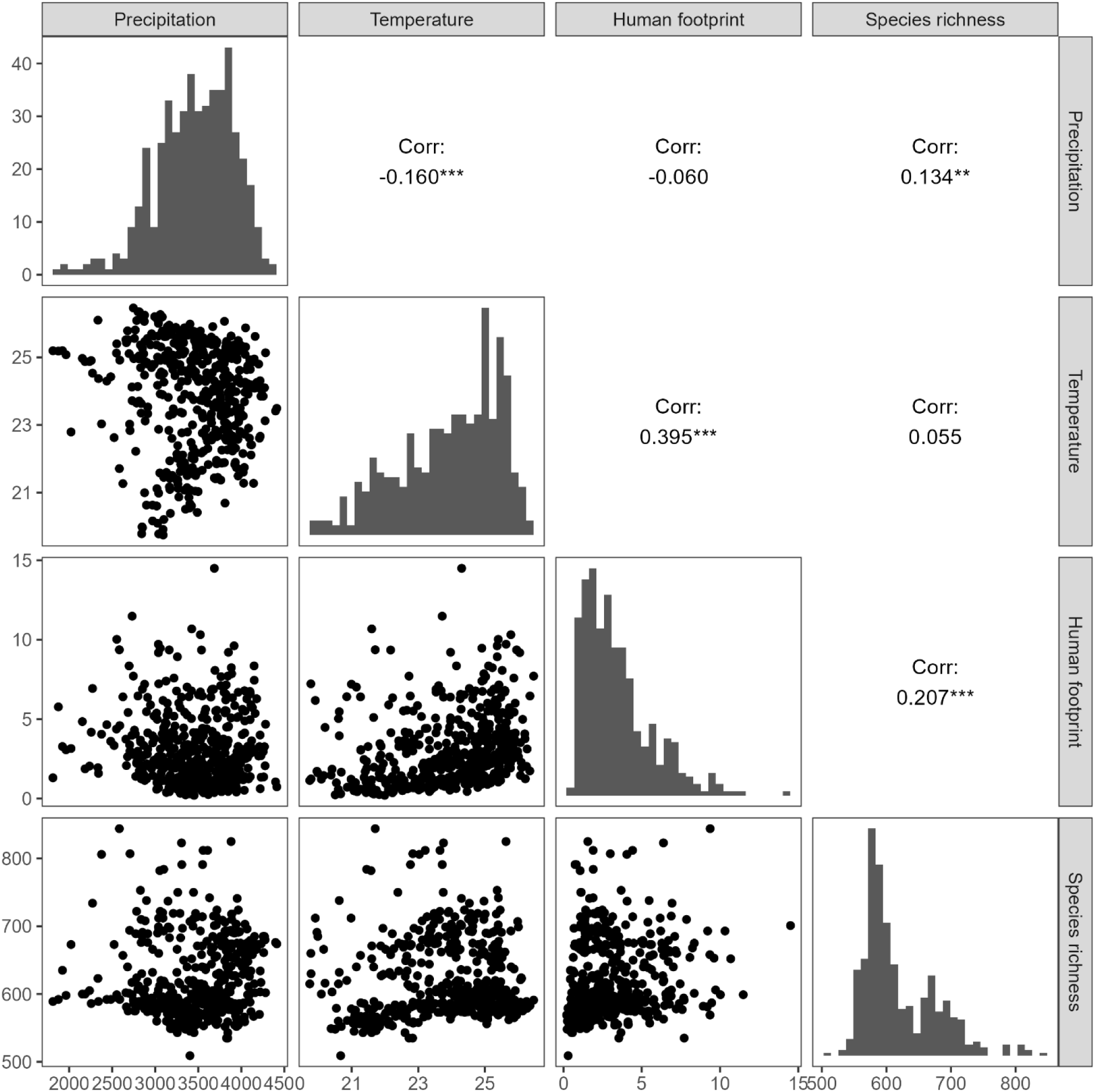
Pairwise correlations between environmental factors. The upper right panels display Spearman’s correlation coefficients with significance levels (***: *p* ≤ 0.001, **: *p* ≤ 0.01, *: *p* ≤ 0.05). The diagonal panels show histograms of each factor’s distribution, while the bottom left panels present scatter plots visualizing pairwise trends.

**Fig. S8.**
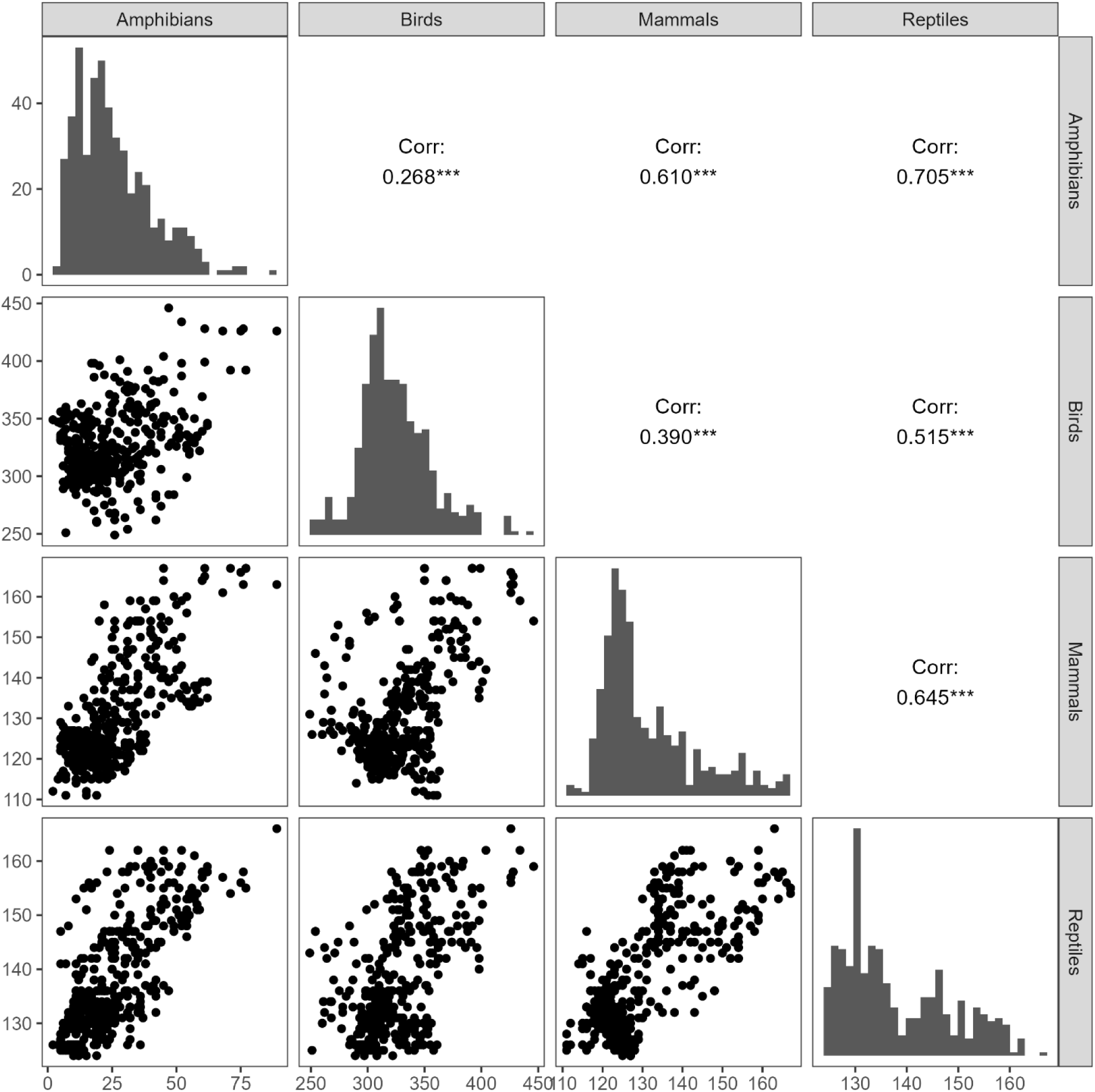
Pairwise correlations between taxa. The upper right panels display Spearman’s correlation coefficients with significance levels (***: *p* ≤ 0.001, **: *p* ≤ 0.01, *: *p* ≤ 0.05). The diagonal panels show histograms of each taxa’s distribution, while the bottom left panels present scatter plots visualizing pairwise trends.

**Fig. S9.**
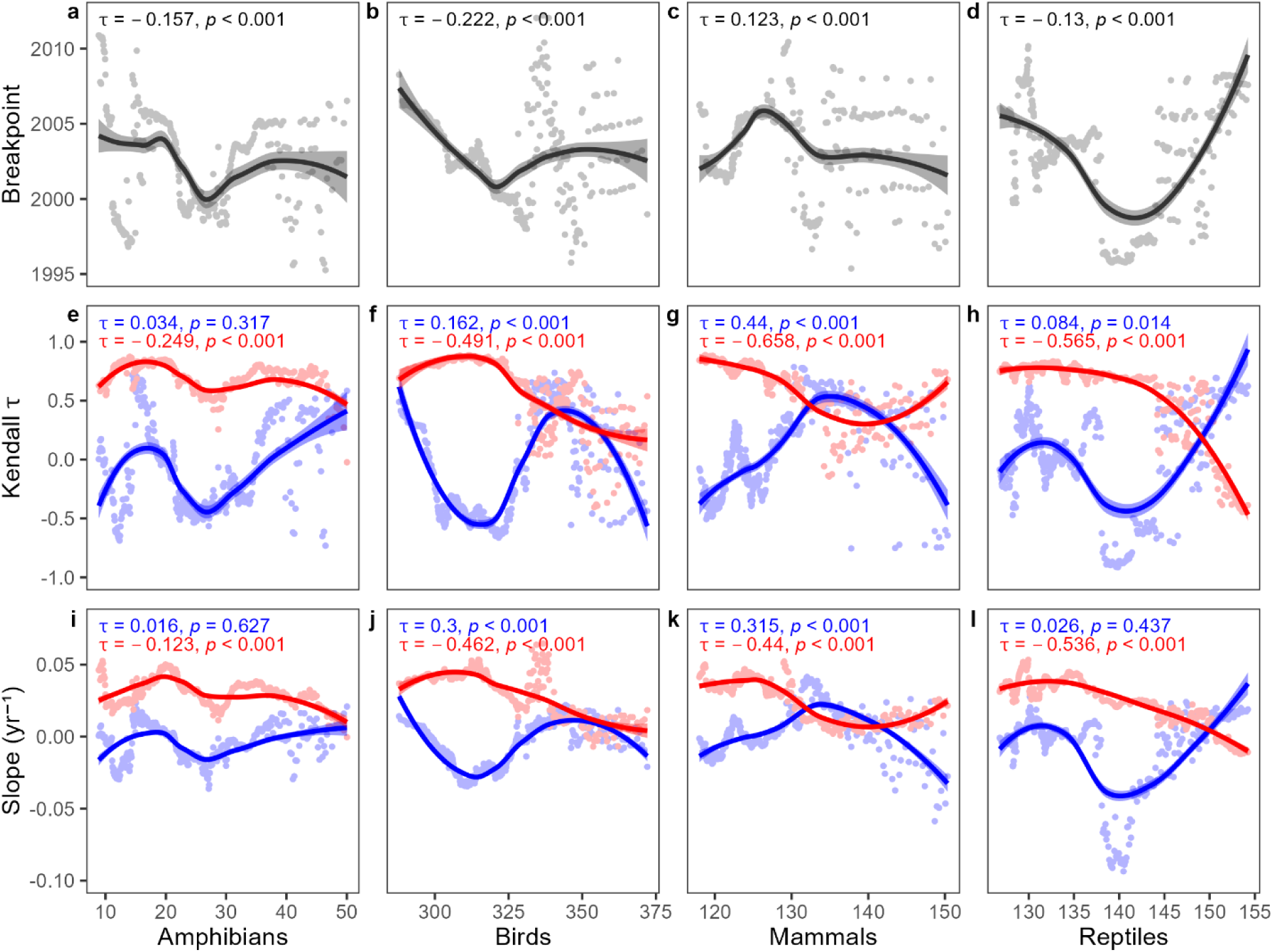
Relationship between vegetation resilience and species richness of different animal taxa (sliding band size = 100 cells). Rows correspond to different resilience metrics and columns represent different factors. **(a, e, i)** Correlation between precipitation and the following resilience metrics calculated on sliding bands of grid cells along a species richness gradient of amphibians: **(a)** posterior mean breakpoint from Bayesian breakpoint regressions; **(e)** Kendall’s *τ* (blue: pre-breakpoint; red: post-breakpoint); and **(i)** posterior mean slope (blue: pre-breakpoint; red: post-breakpoint). The correlations between the resilience metrics and potential influencing factors are estimated using Kendall’s *τ*. the Kendall’s *τ* and associated *p*-values are reported at top of each panel. **(b–d, f–h, j–l)** The same as **(a, e, i)**, but for **(b, f, j)** birds, **(c, g, k)** mammals and **(d, h, l)** reptiles, respectively. In each panel, the values are plotted at the midpoint of each band.

**Fig. S10.**
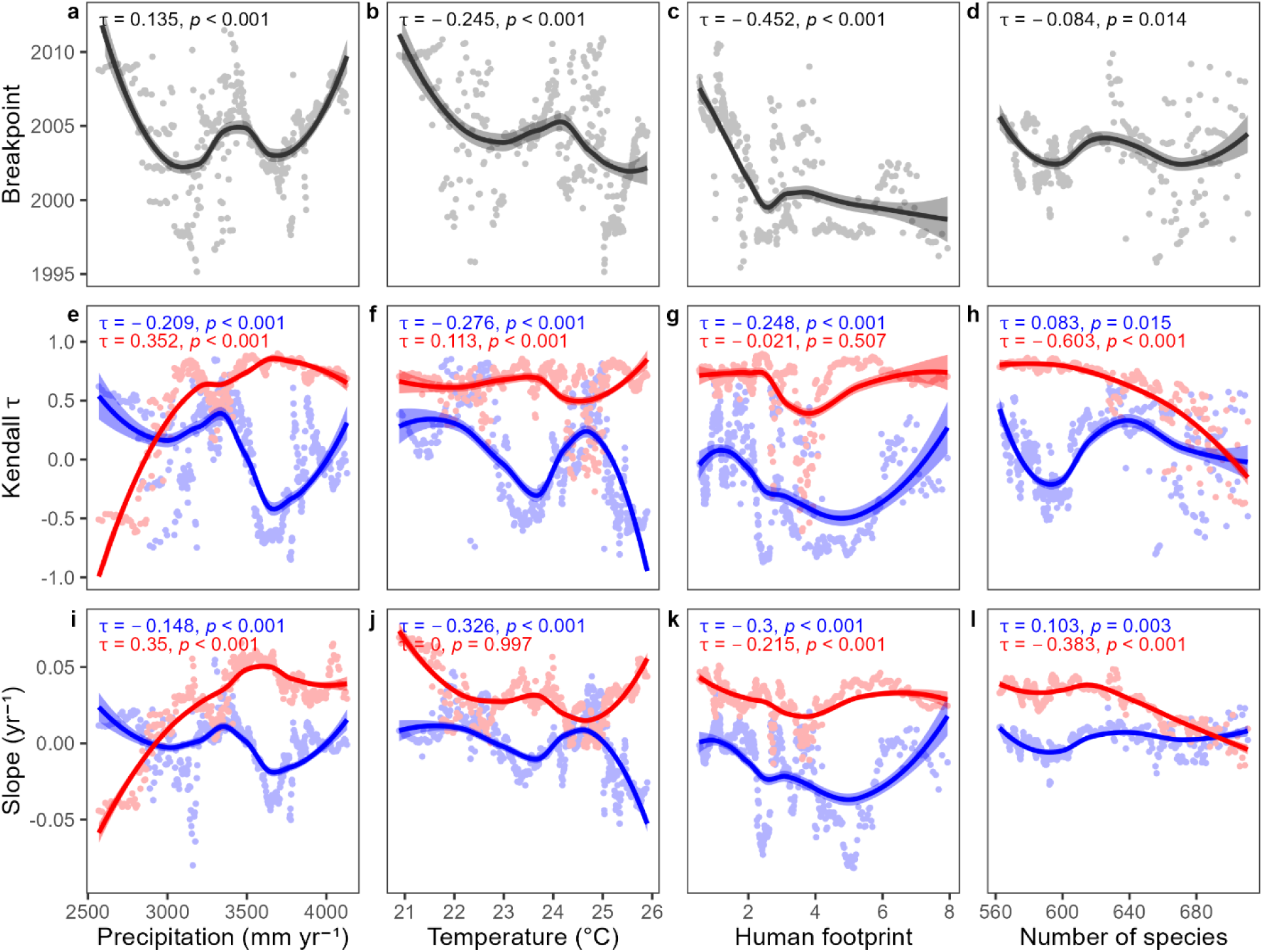
Relationship between potential influencing factors and resilience when varying the band size to 50 cells (recreation of Fig. 3).

**Fig. S11.**
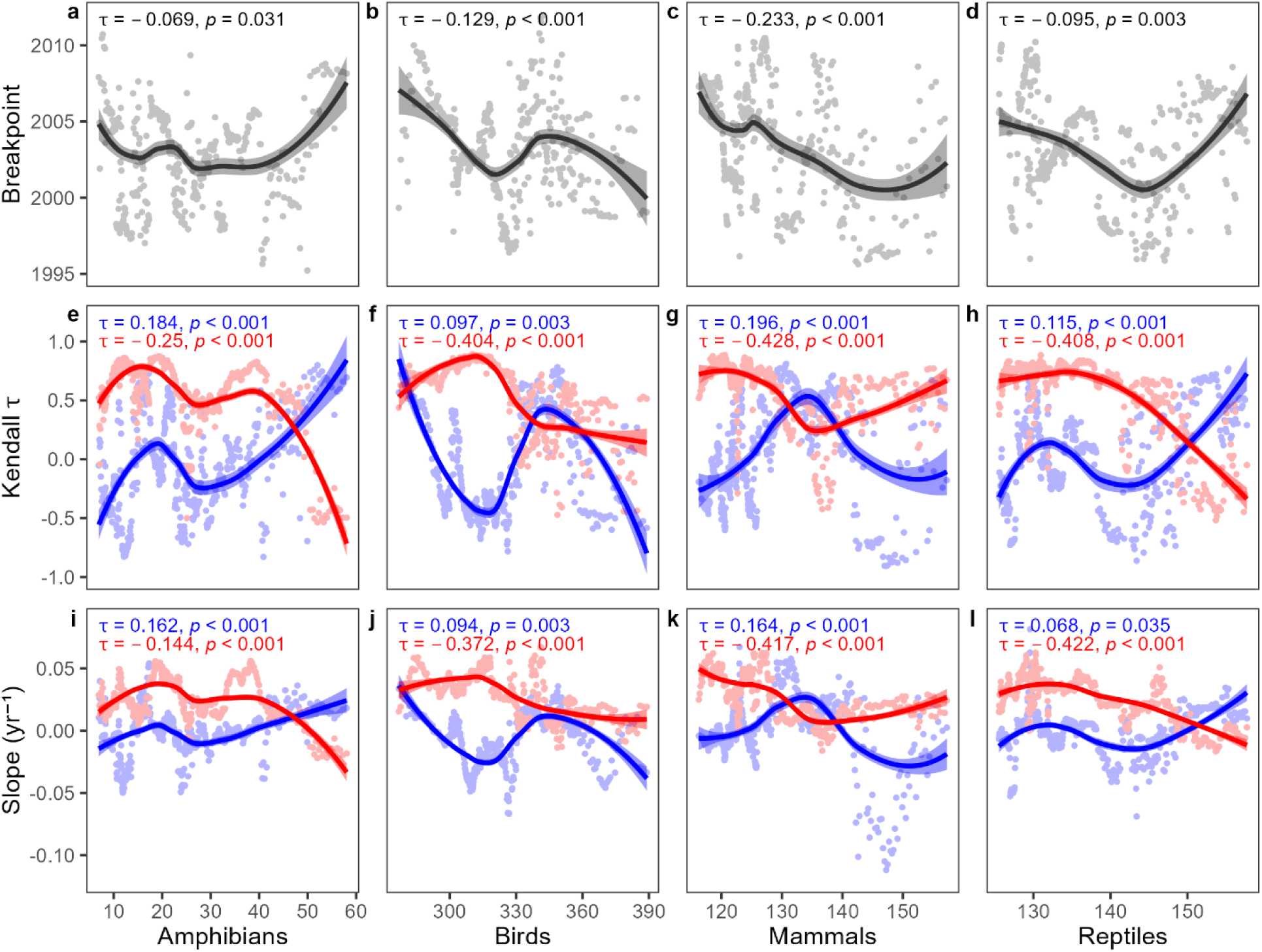
Relationship between species richness of different animal taxa and resilience when varying the band size to 50 cells (recreation of Fig. S9).

## Supplementary Table

**Table S1.** Spearman’s rank correlation coefficients (*ρ*) and associated *p*-values between resilience metrics and periods of individual grid cells (i.e., Figs. 2, S6).

| Region | Correlation between | | Spearman’s $\rho$ | $p$-value |
| --- | --- | --- | --- | --- |
| Borneo | Breakpoint timing | Pre-breakpoint Kendall’s $\tau$ | 0.145 | 0.002 |
|  | Breakpoint timing | Pre-breakpoint slope | 0.135 | 0.003 |
| | Breakpoint timing | Post-breakpoint Kendall’s $\tau$ | 0.084 | 0.007 |
|  | Breakpoint timing | Post-breakpoint slope | 0.150 | 0.001 |
| | Pre-breakpoint Kendall’s $\tau$ | Post-breakpoint Kendall’s $\tau$ | −0.587 | < 0.001 |
|  | Pre-breakpoint slope | Post-breakpoint slope | −0.586 | < 0.001 |
| | Pre-breakpoint Kendall’s $\tau$ | Pre-breakpoint slope | 0.923 | < 0.001 |
| | Post-breakpoint Kendall’s $\tau$ | Post-breakpoint slope | 0.914 | < 0.001 |
| Amazon | Breakpoint timing | Pre-breakpoint Kendall’s $\tau$ | 0.416 | < 0.001 |
|  | Breakpoint timing | Pre-breakpoint slope | 0.428 | < 0.001 |
| | Breakpoint timing | Post-breakpoint Kendall’s $\tau$ | −0.106 | < 0.001 |
|  | Breakpoint timing | Post-breakpoint slope | 0.051 | < 0.001 |
| | Pre-breakpoint Kendall’s $\tau$ | Post-breakpoint Kendall’s $\tau$ | −0.603 | < 0.001 |
|  | Pre-breakpoint slope | Post-breakpoint slope | −0.532 | < 0.001 |
| | Pre-breakpoint Kendall’s $\tau$ | Pre-breakpoint slope | 0.924 | < 0.001 |
| | Post-breakpoint Kendall’s $\tau$ | Post-breakpoint slope | 0.868 | < 0.001 |

**Table S2.** Spearman’s rank correlation coefficients (*ρ*) and associated *p*-values between pre- and post-breakpoint values of Kendall’s *τ* and slope for the spatially-averaged VOD AC(1) of sliding bands of grid cells advancing along gradients of various factors (i.e., between blue and red dots in Figs. 3, S9, S10, S11).

| Factors | Kendall's $\tau$ | | Slope | |
| --- | --- | --- | --- | --- |
| | Spearman's $\rho$ | $p$-value | Spearman's $\rho$ | $p$-value |
| ***Correlations for main factors using sliding bands of 100 grid cells (Fig. 3)*** |  |  |  |  |
| Precipitation | -0.749 | < 0.001 | -0.550 | < 0.001 |
| Temperature | -0.717 | < 0.001 | -0.683 | < 0.001 |
| Human Footprint | -0.496 | < 0.001 | -0.151 | 0.003 |
| Biodiversity | 0.250 | < 0.001 | 0.060 | 0.237 |
| ***Correlations for individual taxa using sliding bands of 100 grid cells (Fig. S9)*** |  |  |  |  |
| Amphibians | 0.269 | < 0.001 | 0.446 | < 0.001 |
| Birds | -0.716 | < 0.001 | -0.556 | < 0.001 |
| Mammals | -0.820 | < 0.001 | -0.434 | < 0.001 |
| Reptiles | -0.241 | < 0.001 | -0.081 | 0.111 |
| ***Correlations for main factors using sliding bands of 50 grid cells (Fig. S10)*** |  |  |  |  |
| Precipitation | -0.577 | < 0.001 | -0.341 | < 0.001 |
| Temperature | -0.583 | < 0.001 | -0.586 | < 0.001 |
| Human Footprint | -0.518 | < 0.001 | -0.084 | 0.081 |
| Biodiversity | -0.460 | < 0.001 | -0.263 | < 0.001 |
| ***Correlations for individual taxa using sliding bands of 50 grid cells (Fig. S11)*** |  |  |  |  |
| Amphibians | -0.124 | 0.010 | -0.179 | < 0.001 |
| Birds | -0.606 | < 0.001 | -0.363 | < 0.001 |
| Mammals | -0.587 | < 0.001 | -0.451 | < 0.001 |
| Reptiles | -0.346 | < 0.001 | -0.170 | < 0.001 |

